# RNA G-quadruplex structure contributes to cold adaptation in plants

**DOI:** 10.1101/2022.03.04.482910

**Authors:** Xiaofei Yang, Haopeng Yu, Susan Duncan, Yueying Zhang, Jitender Cheema, J. Benjamin Miller, Jie Zhang, Chun Kit Kwok, Huakun Zhang, Yiliang Ding

## Abstract

Nucleotide composition is suggested to infer gene functionality and ecological adaptation of species to distinct environments. However, the underlying biological function of nucleotide composition dictating environmental adaptations is largely unknown. Here, we systematically analyzed the nucleotide compositions of transcriptomes across 1000 plants (1KP) and their corresponding habitats. Intriguingly, we found that plants growing in cold climates have guanine (G)-enriched transcriptomes, which are prone to forming RNA G-quadruplex structures. Both immunofluorescence detection and *in vivo* structure profiling revealed that RNA G-quadruplex formation in plants was globally enhanced in response to cold. Cold-responsive RNA G-quadruplexes strongly enhanced mRNA stability, rather than affecting translation. Disruption of individual RNA G-quadruplex promoted mRNA decay in the cold, leading to impaired plant cold response. Therefore, we propose that plants adopted RNA G-quadruplex structure as a molecular marker to facilitate their adaptation to the cold during evolution.

## Main Text

The earth’s prodigious biodiversity was established in part by adaptation to diverse ecological habitats, driving speciation (*1*). In particular, plants are highly evolved to their specific environments, partly due to their sessile nature, with diversity across environments enabling their global colonization (*2, 3*). Varied nucleotide compositions were suggested to affect plant adaptations to specific habitats(*4, 5*). However, understanding of the underlying molecular mechanisms remains unclear.

The recent generation of transcriptome sequences of over 1000 plants (The 1000 plants initiative, 1KP) allows us to systematically assess the biological significance of nucleotide compositions across the plant kingdom (*6*). We analyzed the frequency of the four nucleotides (A: adenine, U: uracil, C: cytosine and G: guanine) over all the major clades of land plants, including dicots, monocots, gymnosperms, ferns, lycophytes and bryophytes (Fig. 1A and Table S1). Overall, the frequencies of both G and C were consistently lower than those of A and U (Fig. 1A). Frequencies among four nucleotides varied much more in the 3’UTR compared to 5’UTR and CDS (Fig. 1A). We then obtained habitat locations of these plants from the Global Biodiversity Information Facility (GBIF) and extracted the corresponding 19 associated bioclimatic variables for these habitats from the WorldClim database (fig. S1A and Table S2, see Methods) (*7*). We calculated the Pearson Correlation Coefficient (PCC) between the transcriptomic nucleotide frequencies and bioclimatic variables primarily concerned with temperature and precipitation (i.e. BIO1: annual mean temperature, BIO12: annual precipitation; full list in Methods). PCCs between nucleotide frequency and temperature bioclimatic variables (BIO1-BIO11) were overall stronger than those between nucleotide frequency and precipitation bioclimatic variables (BIO12-BIO19, fig. S1A), suggesting a stronger relationship between transcriptomic nucleotide composition and temperature of plant habitats. A and U frequencies are mostly positively correlated with temperature bioclimatic variables, while G and C compositions are mostly negatively correlated (fig. S1B). The profound negative correlation between G frequency and temperature bioclimatic variables was notable, indicating an enrichment of G in species growing in cold climates (fig. S1B).

**Fig. 1.**
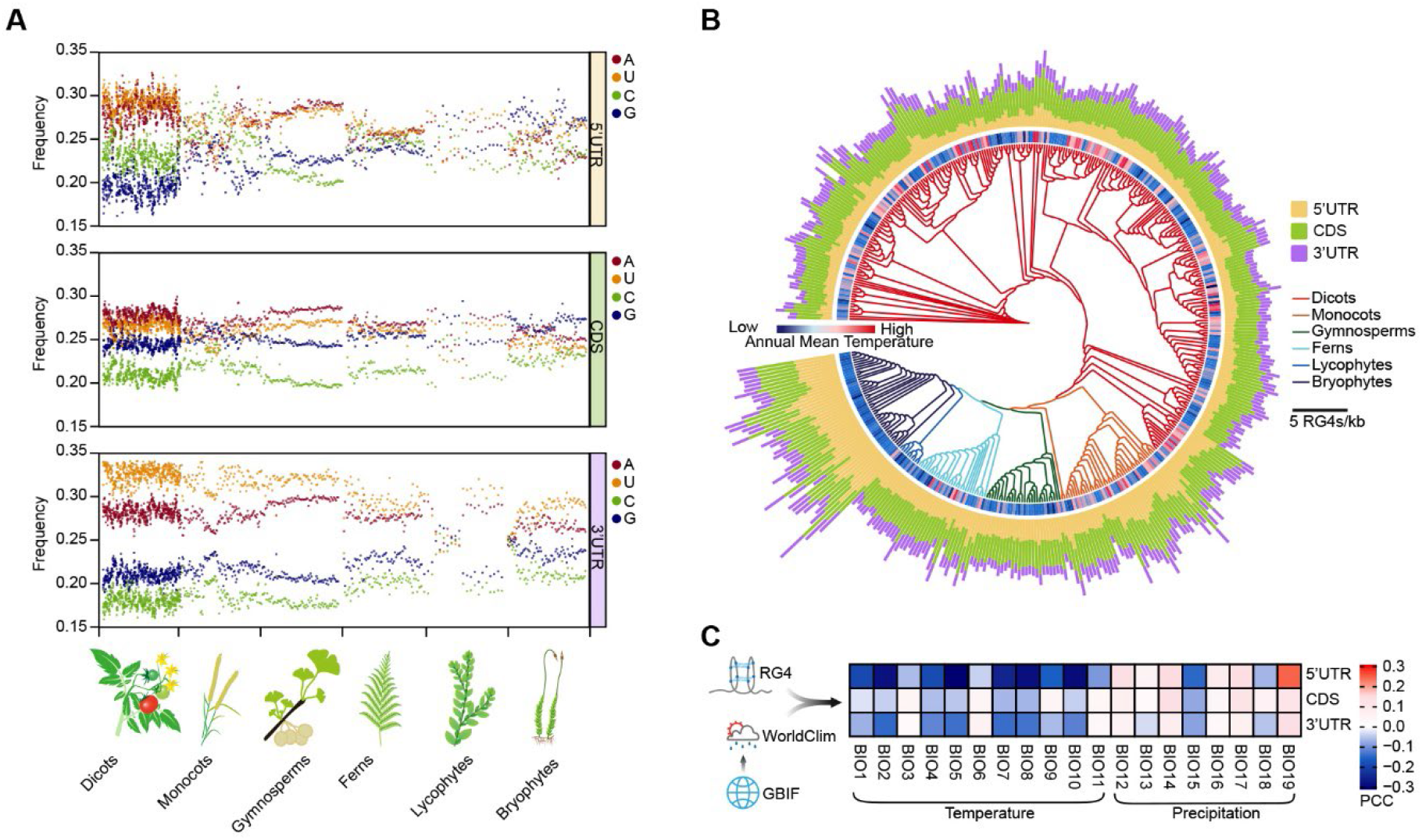
Comparison of transcriptomic guanine (G) frequencies and RNA G-quadruplex frequencies exhibit climatic signatures of plant adaptation. **(A)** Scatter plot showing the frequency of each nucleotide (A: adenine, U: uracil, C: cytosine and G: guanine) in transcriptomes of 906 land plants from the 1000 Plants (1KP Initiative) (*6*). n = 556, 107, 80, 71, 21, 71 for dicots, monocots, gymnosperms, ferns, lycophytes and bryophytes, respectively. **(B)** RNA G-quadruplex (RG4) frequencies in major clades of land plants. Land plants with over 100 occurrences in Global Biodiversity Information Facility (GBIF) (*7*) were included (see Methods). n = 277, 43, 30, 30, 10, 43 for dicots, monocots, gymnosperms, ferns, lycophytes and bryophytes, respectively. The corresponding annual mean temperatures for habitats of plant species were colour-coded. **(C)** Heat plot showing the Pearson Correlation Coefficients (PCCs) between RG4 frequency in plant transcriptomes and associated bioclimatic variables related to plant habitats. PCCs with a *P*-value less than a threshold of 0.01 were retained. Plant habitats were sourced from the GBIF, while associated bioclimatic variables were derived from the WorldClim database (full details see Materials and Methods). Bioclimatic variable names are listed in Methods, such as BIO1: annual mean temperature; BIO12: annual precipitation etc., as explained by WorldClim.

Given that a G-rich region in an RNA molecule is capable of folding into a tertiary RNA structure called RNA G-quadruplex (RG4), involving both Hoogsteen and Watson-Crick base pairs (*8*), we therefore hypothesized that G-enriched transcriptomes from plants prevalent in colder climates may be complemented with RG4 motifs. We then calculated the RG4 frequency across the 1KP dataset, with a median frequency value of 1.27, 2.35 and 0.99 RG4s/kb of RG4 for 5’ UTR, CDS and 3’ UTR, respectively (Fig. 1B and Table S3). Across all genic regions, the overall correlations between RG4 frequency and temperature bioclimatic variables were significantly negative, whist generally less significant correlations between RG4 frequency and precipitation bioclimatic variables were observed (Fig. 1B and 1C), suggesting that RG4 is more strongly enriched in plant species growing in colder climates.

To understand the underlying mechanisms of higher RG4 enrichment in plant adaptation to cold, we first visualized RG4 in cells from the model plant *Arabidopsis thaliana*, using the BG4 antibody, which is able to detect RNA G-quadruplex in cells (*9*). We found a significant increase and enhancement of cytosolic BG4 foci in the cold at 4°C, compared to the normal temperature control at 22°C (Fig. 2A, *P* = 10^−4^, by Student’s t-test). The increase of the BG4 foci in the cold mimicked RG4 stabilization using the well-characterized pyridostatin ligand (PDS) (Fig. 2A, *P* = 10^−6^ by Student’s t-test) (*10*), indicating that cold promotes RG4 folding in plant cells. Notably, when cold-treated plants were returned to 22°C for two hours, BG4 signals recovered to a similar level to that before cold treatment (Fig. 2A, *P* = 0.17 by Student’s t-test).

**Fig. 2.**
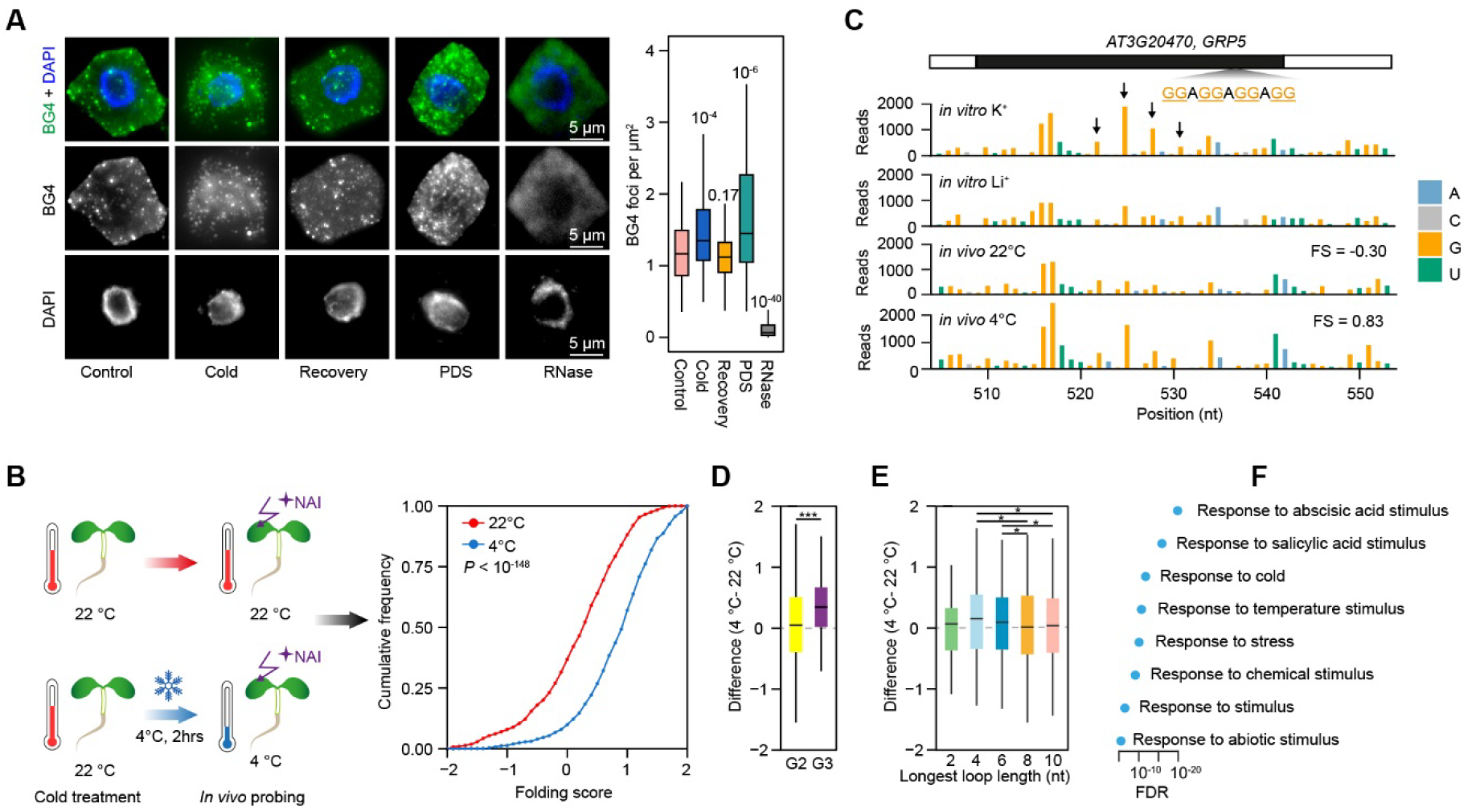
Cold promotes RNA G-quadruplex folding in *Arabidopsis*. **(A)** Immunofluorescence detection of G-quadruplex (RG4) with BG4 antibody in *Arabidopsis* under different thermal conditions. *Arabidopsis* seedlings were grown at 22°C (Control) and were i) treated at 4°C for 2 hours (Cold); or ii) returned to 22°C for 2 hours after cold treatment (Recovery); or iii) treated with 10 μM pyridostatin (PDS); or iv) treated with RNase cocktail (RNase). DAPI indicates the nucleus signal. Statistical analysis was performed using more than 80 cells from 3 individual seedlings for each treatment, significance tested by Student’s t-test in comparison to Control. **(B)** Comparison of RG4 folding by SHALiPE-seq in plants grown at 22°C and treated either at 4°C or 22°C, respectively, n = 817, *P* < 10^−148^, by Student’s t-test. **(C)** SHALiPE-seq profiles of the RG4s on *AT3G20470* (*GRP5*). High reads on last Gs (dark arrow) are compared between *in vitro* benchmarks: either with K^+^ (RG4 folded) or Li^+^ (RG4 unfolded). *In vivo* profiles show unfolded state at 22°C (resembles benchmark with Li^+^) but folded state at 4°C (resembles that with K^+^). **(D)** Frequency of distinct RG4 motifs comparing folding score differences (4°C-22°C) between G2-RG4 and G3-RG4 motifs; n = 1436, 51 for G2 and G3 sites, respectively, \*\*\**P* = 0.006 by Student’s t-test. **(E)** Comparison of RG4 folding score difference frequencies (4°C-22°C) with different RG4 loop lengths. RG4 sites n = 104, 304, 243, 442, and 343 for loop length of 2nt, 4nt, 6nt, 8nt and 10nt, respectively. * *P* < 0.05 by Student’s t-test. **(F)** Representative gene ontology items of transcripts with cold-responsive RG4s.

We then determined the *in vivo* folding status of individual RG4-motifs in the cold using SHALiPE-seq, a high throughput method for RG4 detection (*11*). SHALiPE-seq is based on the preferable modification of the last G residues of G-tracts in the folded RG4 by 2-methylnicotinic acid imidazolide (NAI) (fig. S2) (*11–14*), and therefore detects folded RG4 directly. *Arabidopsis* seedlings grown at 22°C were treated for 2 hours either at 4°C or 22°C (control) followed by treatment with NAI at 4°C or 22°C, respectively, before generating SHALiPE-seq libraries for both groups (fig. S2). Comparable NAI modification efficiencies under different temperatures were achieved, as illustrated (figs. S3 and S4), by both gel-based analysis and SHALiPE-seq library analysis on the 18S rRNA.

Reads distribution for Gs of SHALiPE-seq were characterized using the Gini index, where a high Gini index indicates an uneven distribution of reads on Gs for folded RG4s, whilst a low Gini index indicates an even distribution of unfolded RG4s (*11, 14*). We initially established Gini index benchmarks for folded or unfolded RG4s under *in vitro* conditions with either K^+^ (stabilizes RG4) or Li^+^ (destabilizes RG4) (fig. S2) (*11*). We then implemented a quantitative algorithm to calculate the folding status *in vivo,* which we termed ‘folding score’: a comparison of Gini *in vivo* with Gini *in vitro* in the presence of Li^+^, scaled relative to Gini *in vitro* with K^+^ vs Gini *in vitro* with Li^+^ (see methods) (*11*). The folding scores at 4°C were overall significantly higher compared to those at 22°C (Fig. 2B, *P* <10^−148^, by paired Student’s t-test, Table S4), further supporting that stronger RG4 folding occurs in the cold. In particular, some RG4s showed a switch from unfolded to folded status as the temperature shifted from 22°C to 4°C. *In vivo* SHALiPE-seq profile examples of RG4s on *AT3G20470* at 22°C resembled that of the unfolded *in vitro* benchmark in the presence of Li^+^, while corresponding *in vivo* SHALiPE-seq profiles at 4°C resembled that of the folded *in vitro* benchmark in the presence of K^+^ (Fig. 2C). Therefore, our SHALiPE-seq data along with our immunofluorescence visualizations reinforce the notion that RG4 folding is responsive to cold in living plants.

RG4s have a range of G-quartets and loop lengths (*8, 15*). To test whether different types of RG4s may be favoured in the cold, we compared the folding score differences between 4°C and 22°C on individual RG4s. We found greater folding score differences for RG4s with three G-quartets (G3-RG4) compared to those with two G-quartets (G2-RG4, Fig. 2D, *P* = 0.006, by Student’s t-test). In addition, the folding status of RG4s with intermediate lengths from 4nt to 6nt were more easily enhanced in the cold, compared to RG4s with either a very short loop of 2nt or a very long loop of over 8nt (Fig. 2E, *P* < 0.05, by Student’s t-test). Thus, cold conditions are likely to strongly promote the folding of RG4s with a higher number of G-quartets and intermediate loop lengths.

To understand any biological relevance between RG4s and the cold, we performed gene ontology (GO) analysis on genes containing cold-responsive RG4s with higher folding scores upon cold treatment. We found specific transcript-related enrichment in biological functions, such as response to abiotic stimulus, response to temperature stimulus and response to cold (Fig. 2F). Thus, those genes containing cold-responsive RG4s are also likely to function in plant response to cold.

To further explore how these cold-responsive RG4s may affect the plant response to cold, we assessed their molecular functions in regulating gene expression. Extensive studies have suggested that RG4 serves as a translational repressor (*8, 11, 16*). Thus, we assessed whether cold-responsive RG4s repress translation in the cold. We compared the translation efficiencies (TEs) of those genes containing cold-responsive RG4s at either 4°C and 22°C. Overall, we observed only a subtle variation (~2%) and a high correlation between TE at 4°C and TE at 22°C (Fig. 3A, PCC = 0.86, difference = 2%, Table S5), suggesting cold-responsive RG4s are unlikely to contribute much to translation. We then examined the folding score differences between 4°C and 22°C of RG4s in different genic regions and found RG4s in the 3’UTR possessed the greatest folding score differences (Fig. 3B, *P* = 0.01 for CDS vs 3’UTR, by Student’s t-test). Based on previous studies that suggested structural elements in 3’UTRs are likely to regulate mRNA stability (*17–19*), we hypothesized that cold-responsive RG4s may impact mRNA stability. We measured the RNA stability of the *Arabidopsis* transcriptome at both 22°C and 4°C using the transcription arrest assay (Fig. 3C) (*20*). Upon transcription arrest, the rate of mRNA abundance decline reveals RNA stability, whereby a rapid decrease indicates low stability whilst a slow decrease indicates high stability. Overall, we found a generally slower mRNA abundance decline at 4°C compared to 22°C (Fig. 3D). Notably, the cold effect of slowing down RNA abundance decline was found to be highest for transcripts with cold-responsive RG4s in 3’UTRs, compared to those transcripts without RG4s (non-RG4), or with cold-responsive RG4s in 5’UTR and/or CDS regions (Fig. 3D). The transcription arrest profiles for individual mRNAs are illustrated for *AT1G13390*, *AT4G32020* and *AT5G24930* along with their corresponding qRT-PCR validations (fig. S5).

**Fig. 3.**
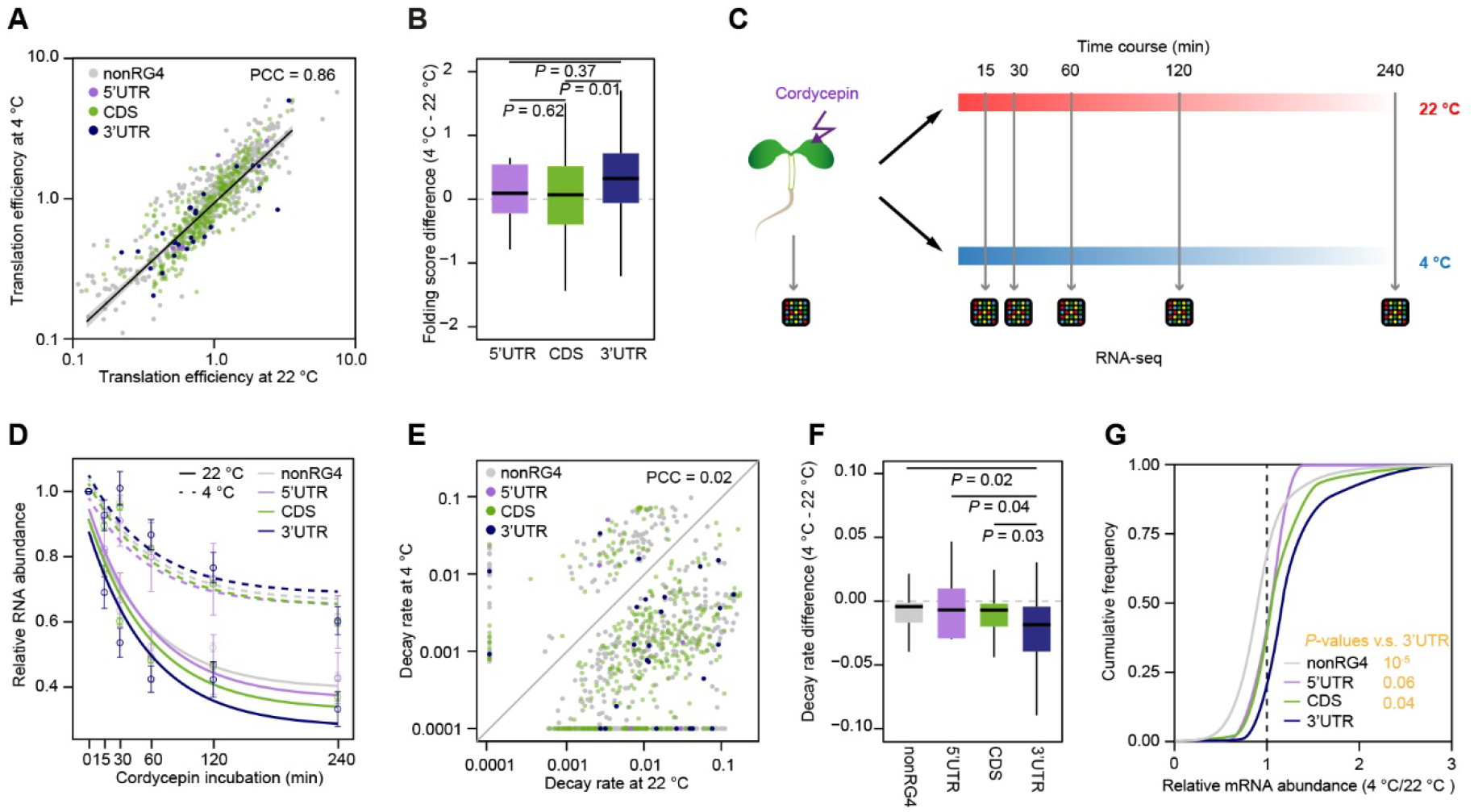
Cold-responsive RNA G-quadruplex in 3’UTR represses mRNA decay. **(A)** Comparison of translation efficiency (TE) at 22°C and 4°C, for transcripts without RG4 (nonRG4), with RG4 in 5’UTR, with RG4 in CDS and with RG4 in 3’UTR. The number of transcripts, n = 465, 6, 531, 36, for nonRG4, 5’UTR, CDS and 3’UTR respectively. **(B)** Comparison of folding score differences (4°C - 22°C) between RG4 located in different genic regions. The number of RG4 sites, n =9, 1421, 57 for 5’UTR, CDS and 3’UTR, respectively; *P*-values by Student’s t-test. **(C)** Schematic of determining RNA stability in *Arabidopsis* at 22°C and 4°C, by transcriptional arrest analysis with cordycepin incubation. **(D)** RNA abundance curves at 22°C and 4°C for transcripts with cold-responsive RG4s, otherwise in Fig. 3A. Error bar indicates SE of different transcripts. **(E)** Comparison of decay rate of transcripts with cold-responsive RG4s at 22°C and 4°C, otherwise in Fig. 3A. **(F)** Comparison of decay rate differences (4°C - 22°C) on transcripts with cold-responsive RG4s. *P*-value by Student’s t-test, otherwise in Fig. 3A. **(G)** Line plot showing relative mRNA abundance at 4°C versus 22°C for transcripts with cold-responsive RG4s. *P*-values by Student’s t-test, otherwise in Fig. 3A.

We then derived the mRNA decay rate for individual mRNAs with cold-responsive RG4s at both 4°C and 22°C (Table S6). The mRNA decay rates at 4°C altered dramatically compared to those at 22°C (Fig. 3E, PCC = 0.02), and generally showed slower mRNA decay rates in the cold. We subsequently calculated the decay rate differences between 4°C and 22°C and found a significantly greater difference of decay rate between 4°C and 22°C on transcripts with cold-responsive RG4s in 3’UTR compared to that of transcripts without RG4s or with RG4s in 5’UTR or CDS regions (Fig. 3F, *P*-values = 0.02, 0.04 and 0.03 respectively, by Student’s t-test). As mRNA stability directly affects steady-state mRNA abundance, we reasoned that cold enhances the steady-state mRNA abundance of transcripts with cold-responsive RG4s. Indeed, we found a strong increase of mRNA abundance in the cold for transcripts with cold-responsive RG4s in the 3’UTRs, rather than nonRG4 transcripts or cold-responsive RG4s in 5’UTR/CDS regions (Fig. 3G, *P* = 10^−5^, 0.06 and 0.04 respectively, by Student’s t-test). Collectively, our results suggest cold-responsive RG4s strongly impact mRNA stability rather than mRNA translation. These cold-responsive RG4s in the 3’UTR serve as mRNA stabilizers, reducing mRNA degradation in the cold.

Considering the strong molecular function of these cold-responsive RG4s in 3’UTRs, we reasoned that these cold-responsive RG4s, as mRNA stabilizers, may affect plant phenotypic response to the cold. Of the cold-responsive RG4s present in 3’UTRs, the one with the greatest folding score difference between 4°C and 22°C was found on *AT1G13390* (fig. S6A and Table S4), designated as *CORG1* (<u>CO</u>ld-responsive <u>R</u>NA <u>G</u>-quadruplex <u>1</u>), with the annotation encoding a translocase subunit protein (courtesy of The Arabidopsis Information Resource). To further explore whether this *CORG1* RG4 contributes to mRNA stabilization in the cold, we constructed the reporter gene, *FIREFLY-LUCIFERASE* (*FLUC*) fused with the 3’UTR of *CORG1* containing the wildtype cold-responsive RG4 (wtRG4) or a disrupted RG4 (mutRG4, Fig. 4A). We then performed transcriptional arrest assays at both 4°C and 22°C to measure the corresponding mRNA stabilities. The *FLUC* mRNA with the 3’UTR of *CORG1* containing the mutRG4 degraded much more rapidly compared to that containing wtRG4 at 4°C, while no significant differences were found between these two versions at 22°C (Fig. 4B), indicating that this cold-responsive RG4 enhanced mRNA stabilization in cold conditions.

**Fig. 4.**
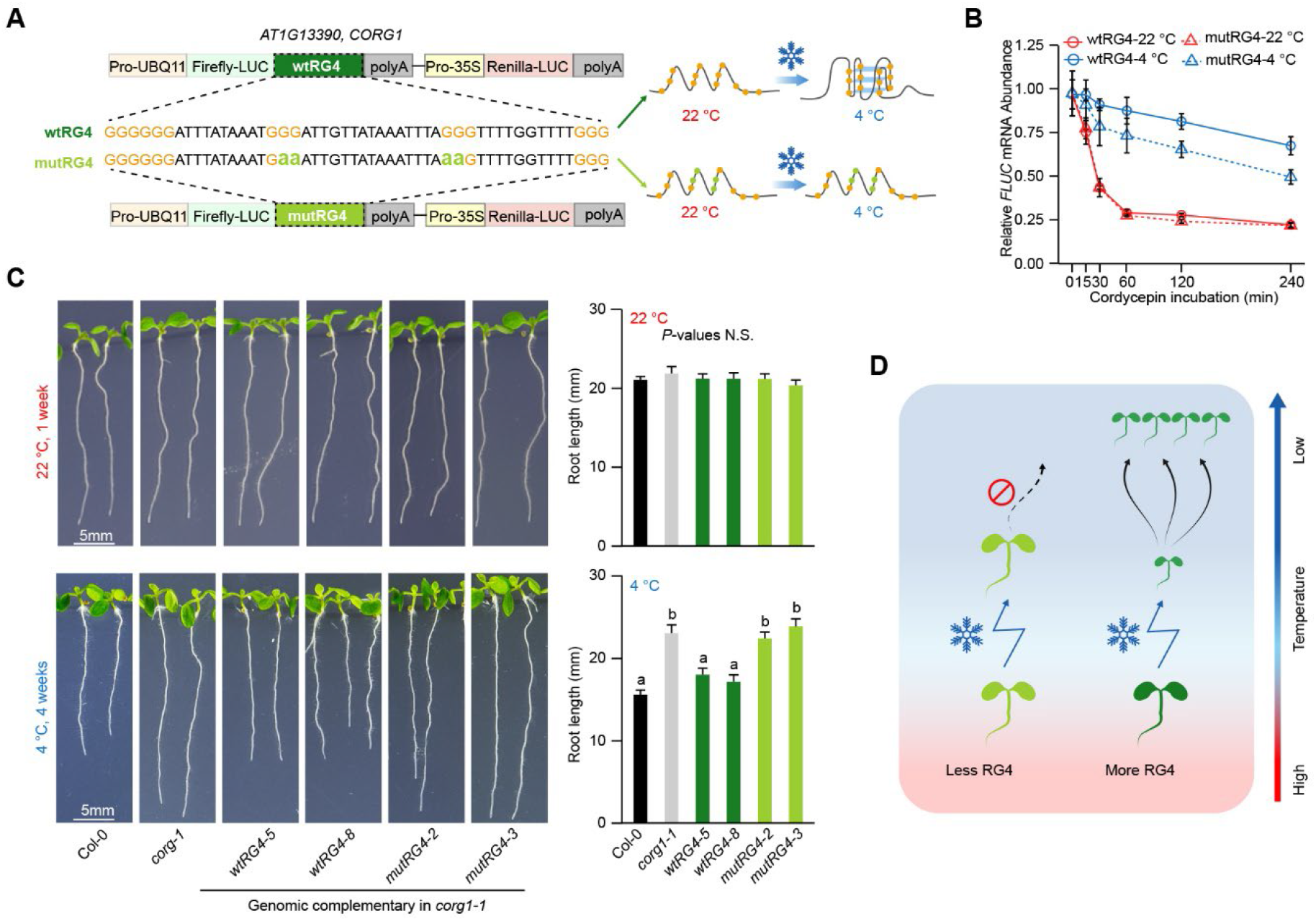
RNA G-quadruplex regulates plant cold-responsive growth. **(A)** Schematic of folding status of wild-type RG4 (wtRG4) and mutated RG4 (mutRG4) on *CORG1* 3’UTR at 22°C and 4°C. The RG4 folding potential was either maintained in wtRG4 or attenuated in mutRG4 by G to A substitution. **(B)** RNA decay curves at 22°C and 4°C for *FIREFLY LUCIFERASE* (*FLUC*) transcripts with wtRG4 or mutRG4, as with the designs shown in Fig. 4A. Error bar indicates SE, n = 4. **(C)** Phenotypes of plants of different genotypes grown at 22°C or 4°C. wtRG4 or mutRG4 denotes *corg1-1* mutant complemented with genomic DNA of *CORG1* carrying wtRG4 or mutRG4 respectively, as illustrated in Fig. 4A. Comparison was performed on the plants grown vertically on plates at 22°C for 1 week, or at 4°C for 4 weeks. Significance tested by one-way ANOVA/Tukey HSD post hoc test (*P* < 0.05), n ≥ 20, error bar indicates SE. **(D)** Model showing the selective enrichment of RG4 driving plant adaptation to cold climates. More RG4s in the plant transcriptome enable higher phenotypic plasticity in cold sensing, therefore shaping plant adaptation to colder climates.

We then reintroduced *Arabidopsis* genomic DNA of *CORG1* containing 3’UTR with either wtRG4 or mutRG4 into the *corg1-1* null mutant (figs. S6, B and C). At 22°C, the primary root growth of *corg1-1* was similar to that of wild type Col-0 (Fig. 4C). At 4°C, the root length of *corg1-1* was distinctively longer than that of Col-0, indicating *CORG1* represses plant growth under cold conditions. When the *corg1-1* mutant was complemented with *mutRG4-CORG1,* the root length was significantly longer than *wtRG4-CORG1* complemented plants at 4°C, while no significant differences were observed at 22°C (Fig. 4C, *P*-values < 0.01, one-way ANOVA/Tukey HSD post hoc test), suggesting impairment of *mutRG4-CORG1* plants in sensing cold. The cold sensitive defects in the *mutRG4-CORG1* plants correspond to the more rapid RNA decay and decrease in steady-state *CORG1* RNA abundance (figs. S6, D and E). Collectively, these results indicate this cold-responsive RG4 regulates *CORG1* mRNA stability in the cold and so significantly modulates plant cold-responsive growth.

Our broad and deep investigations of nucleotide composition across the plant kingdom promote the discovery of RNA G-quadruplex structure serving as a molecular marker shaping plant environmental adaptation. RNA G-quadruplexes embedded across the plant transcriptome globally facilitate stabilizing RNA molecules in response to cold (Fig. 4D). Hence, our results suggest that RNA structure, as a key regulator of gene expression (*21–23*), may have evolved due to selection pressures arising from distinct temperature environments. Our work therefore advances fundamental understanding of the molecular mechanisms underlying species adaptation, translation of this conceptual advance to molecular engineering strategies could help to address global climate change challenges indicated by continuing increases in the frequency, intensity and duration of extreme low-temperature events (*24, 25*).

## Acknowledgments

We thank Prof. Beverley Glover (U. Cambridge), Dr. Sam Brockington (U. Cambridge), Prof. Shiheng Tao (Northwest A&F University), Prof. Giles Oldroyd (SLCU, Cambridge) and Dame Prof. Caroline Dean (John Innes Centre), Dr. Antony Dodd (John Innes Centre), Prof. Cristobal Uauy (John Innes Centre), Prof. Alison Smith (John Innes Centre) and Dr. Desmond Bradley (John Innes Centre) for discussions with this work.

## Author contributions

Y.D. conceived the study; X.Y., H.Y., J.B.M., C.K.K., H.Z., and Y.D. designed the study; X.Y., S.D., Y.Z., and J.Z. performed the experiments; H.Y., X.Y., S.D. and J.C. did the analyses; H.Z., and Y.D. supervised the analyses; X.Y., H.Y., S.D., J.C., H.Z., and Y.D. wrote the paper with input from all authors.

## Data and materials availability

The raw data have been deposited in the Sequence Read Archive (SRA) (https://www.ncbi.nlm.nih.gov/sra) under BioProject ID number PRJNA762705. SHALiPE-seq data *in vitro* and *in vivo* at 22°C are available under BioProject ID number PRJNA561194 (Yang et al., 2020). All data are available in the main text or the supplementary materials.

## Supplementary Materials for

**Other Supplementary Materials for this manuscript include the following:**

Captains for Table S1 to Table S7

Table S1: Nucleotide frequency of transcriptomes of land plants in One Thousand Plant Transcriptomes Initiative (1KP)

Table S2: Geographic and bioclimatic features for land plants in One Thousand Plant Transcriptomes Initiative (1KP)

Table S3: RG4 density in land plants in One Thousand Plant Transcriptomes Initiative (1KP)

Table S4: Folding score of RG4s at 22°C and 4°C in *Arabidopsis*

Table S5: RNA-seq and Polysome-seq at 22°C and 4°C in *Arabidopsis*

Table S6: RNA decay rate at 22°C and 4°C in *Arabidopsis*

Table S7: Plant genotypes, primers used in this study

## Materials and Methods

### Plants and growth conditions

The *Arabidopsis thaliana* ecotype columbia (Col-0) and *corg1-1* mutant (SALK_097604) with T-DNA insertion were obtained from the Nottingham Arabidopsis Stock Centre (NASC); the homozygous mutant was identified using PCR-based genotyping (*1*). *Arabidopsis* seeds were sterilized using 70% ethanol for 10 min, washed with distilled water for 3 times and plated on half-strength Murashige and Skoog medium supplemented with 1% sucrose. After standing at 4°C for 3 days, the plates were placed in a growth chamber at 22°C. To complement the mutant, *corg1-1* mutant plants were transformed using a floral dipping method (*2*). Transgenic plants were selected using GM media with 10 mg/l phosphinotricin.

### Plasmid construction

To complement the *Arabidopsis* mutants with wild-type sequence, a DNA fragment of *AT1G13390* was amplified from Col-0 genomic DNA using CloneAmp HiFi PCR Premix (Clontech) with primers listed in Table S7. To complement the *Arabidopsis* mutants with disrupted RG4 sequence (mutRG4), fragments were amplified by overlap PCR with designed mutations (guanine −> adenine, G −> A) on primers (Table S7) (*3*). PCR products were further introduced into the XmnI and KpnI digested pB7FWG2.0 with In-Fusion (Clontech). For dual-luciferase analysis, the sequence of 3′UTR of *AT1G13390* with or without mutation was cloned into the expression vector inter2 digested with AscI and PstI as introduced with designed primers (Table S7). Sequencing-confirmed vectors were then transformed to *Agrobacterium tumefaciens* GV3101.

### Immunofluorescence detection of RG4

*Arabidopsis* seedlings of 5 days after germination (DAG) were fixed in 4% methanol-free formaldehyde (Thermo Scientific) diluted in nuclease free 1 x PBS (Invitrogen) for 30 minutes, then washed 3 times with 1×PBS (Invitrogen). Roots were then squashed under coverslips onto slides that had been pre-treated with 3-Aminopropylthiethoxysilane (Sigma-Aldrich) to increase tissue adherence (*4*). Samples were then immersed in liquid nitrogen for around 10 seconds before the coverslips were removed. The tissue was left to dry for 1hr at room temperature, 1×PBS with 0.1% Triton X-100 was added to each slide, incubated for 3 mins following with three washes with 1×PBS. At this stage RNase treatment was carried out for a subset of +RNase slides by adding 12U RNase A and 200U RNase T1 (Invitrogen) diluted in 1×PBS to each slide, and incubated for 1.5 hours at 37 °C following with three 5-min washes with 1×PBS. All slides were immersed in 70% ethanol for 2 hours at room temperature. After three 1×PBS washes, 2% BSA dissolved in 1×PBS was added to each slide and left to incubate at room temperature for 1 hour. The block was aspirated and anti-DNA/RNA G-quadruplex Mouse Fab fragment antibody (BG4 antibody, His-Tagged, Lambda antibody AB00174-1.6, Absolute Antibody) with a dilution of 1:300 in 1× PBS with 2% BSA was added to each slide and incubated in a dark (*5*), humid box overnight at 4°C. Three 5-min washes with 1×PBS were then carried out and 1:1000 diluted Alexa Fluor 647 goat anti-mouse IgG H+L secondary antibody A21235 (Invitrogen) was added to each slide, followed by incubation at room temperature for 1 hour. Two 5-min washes with 1× PBS were carried out followed by a 30 min incubation at room temperature with 1 μg/mL DAPI in 1×PBS, following with three 5-min washes with 1× PBS and three washes with nuclease-free water. The samples were mounted with ProLong Glass (Thermo Fisher Scientific) under No. 1.5 coverslips (VWR). Root cells were imaged using a 100x objective (NA 1.46) on an inverted ZEISS Elyra PS1 microscope (ZEISS, Oberkochen) equipped with an Andor EM-CCD camera and controlled by ZEN 2.3 SP1 software (ZEISS, Oberkochen). 0.2 μm optical Z-sections were carried out and images were obtained using 405 nm and 642 nm laser lines. ImageJ plugin RS-FISH was used for automated G-quadruplex quantification as described (*6, 7*). Analysis was carried out on 2D maximum intensity Z-projections for individual cells with default settings apart from optimization of difference-of-gaussian values and intensity threshold. To perform PDS treatment in *Arabidopsis*, seedlings were incubated with 10 μM pyridostatin (PDS) dissolved in half strength of Murashige and Skoog liquid medium for 2 hours. PDS penetration was carried out by vacuum infiltration - three times for 30 seconds and 1 minute break each. To perform cold treatment on *Arabidopsis*, the plate was placed into growth chamber MIR-254-PE (Panasonic) at 4°C for 2 hours.

### *In vivo* NAI probing of RNA

The NAI probing buffer was kept in an incubator at the desired temperature overnight before use. NAI probing at 22°C was carried out as described in our previous study (*8*). For NAI probing at 4°C, plates with 5-DAG etiolated seedlings were treated at 4°C for 2 hours. Treated seedlings were harvested in a cold room and incubated with 400 mM NAI at 4°C for 15 minutes. NAI was quenched using 5 times of DTT to that of NAI. Seedlings were washed 3 times with distilled water, ground in liquid nitrogen and applied to RNA extraction using RNeasy Plant Mini Kit (Qiagen).

### Gel-based analysis of NAI probing

RNA was extracted after NAI probing. 1 μg of total RNA was dissolved in 6 μL water, 1 μL of 5 μM Cy5 modified RT primer for 18S rRNA (listed in Table S7) and 0.5 μL of 10 mM dNTPs were denatured at 95°C for 3 min. The reaction was cooled down to 50°C before adding 2 μL of 5× home-made RT buffer (100 mM Tris (pH 8.3), 500 mM LiCl, 15 mM MgCl2, 5 mM DTT) and 0.5 μL reverse transcriptase Superscript III (Invitrogen) and mixing quickly with a pipette. The RT reaction was incubated at 50 °C for 20 min following a staying at 85 °C for 10 min to inactivate reverse transcriptase. cDNA hybridized RNA was degraded by adding 0.5 μL of 2M NaOH and incubating at 95°C for 10 min. Equal volumes of 2X stopping dye (95% formaldehyde, 20 mM EDTA (pH8.0), 20 mM Tris (pH 7.5), orange G) were added and incubated at 95°C for 5 min. The resultant reaction was kept at 65°C and loaded to 8% Acrylamide:Bis-Acrylamide-Urea gel for electrophoresis. For the sequencing lanes, RNA was dissolved in 5 μL water, 1 μL 10mM of corresponding ddNTP (Roche) was added at the beginning.

### Generation of the SHALiPE-seq libraries

Libraries of SHALiPE-seq were prepared as described (*8, 9*). PolyA-selected RNA was recovered and reverse transcribed using superscript III (Invitrogen) and RT primer (5’CAGACGTGTGCTCTTCCGATCTNNNNNN3’) with home-made RT buffer (20 mM Tris (pH 8.3), 100 mM LiCl, 3 mM MgCl_2_, 1 mM DTT). The 3’end of resultant cDNAs were ligated to a ssDNA linker (5’-PhosNNNAGATCGGAAGAGCGTCGTGTAG-/3SpC3/3’) using Circligase ssDNA Ligase (Epicentre) at 65°C for 12 hrs. Product longer than 100 nt was recovered using QIAquick Gel Extraction Kit (Qiagen) after separation of the ligation products with TBE-Urea 10% Gel (Invitrogen). Purified cDNA was subjected to PCR amplification using KAPA Library Amplification Kits (Roche) with Forward Library primer (5’AATGATACGGCGACCACCGAGATCTACACTCTTTCCCTACACGACGCTCTTCCGATCT3’) and Reverse Library primers (5’CAAGCAGAAGACGGCATACGAGATNNNNNNGTGACTGGAGTTCAGACGTGTGCTCTTCCGATC3’, where NNNNNN denotes the barcodes, e.g. Index1 is CGTGAT for Illumina sequencing). Three rounds of agarose gel purification were performed to purify the fragments of 200~650 bp using QIAquick Gel Extraction Kit (Qiagen). The libraries were sequenced with Illumina HiSeq 4000 platform by BGI Genomics (Hongkong).

### Polysome associated mRNA analysis and RNA-seq

Polysome associated mRNA analysis was performed as described with modifications (*10, 11*). Briefly, around 500 mg 5-DAG etiolated seedlings (grown at 22°C, with or without 2 hours treatment at 4°C) were harvested and ground into fine powder in liquid nitrogen. The powder was dissolved in 500 μL precooled polysome extraction buffer (200 mM Tris-HCl, pH 8.4, 50 mM KCl, 1% deoxycholic acid, 25 mM MgCl_2_, 2% Polyoxyethylene 10 tridecyl ether, 2 mM DTT, 400 U/mL recombinant Rnasin, 50 μg/mL cycloheximide). After incubation on ice for 30 minutes and centrifugation at 13200 rpm for 15 min at 4°C, 500 μL supernatant was transferred to 15-60% sucrose gradient. Polysomes were fragmented by 4 hours of centrifugation at 40000 rpm in Beckman ST40Ti rotor. Low to high sucrose gradient was collected, taking fractions 1 to 12. Fractions of 7 to 12 representing the translation level were taken for RNA isolation with TRIzol reagent (Ambion). To perform RNA-seq experiment, around 25 mg plant powder was used for direct RNA isolation by RNeasy Plant Mini Kit (Qiagen). Polysome-bound RNA or total RNA was subjected to RNA-seq library generation by BGI Genomics following the manufacturer’s BGISEQ-500 protocol.

### Transcriptional arrest

Transcriptional arrest analysis was performed as described with modifications (*12*). For analysis in *Arabidopsis*, seedlings were harvested, and placed into a petri dish with two layers of filter paper and incubation buffer with cordycepin (1mM PIPES, pH 6.25, 15 mM sucrose, 1 mM potassium chloride, 1 mM sodium citrate and 1mM cordycepin). For reporter assay in tobacco *N. benthamiana*, leaves after 48 hours of agroinfiltration were harvested, and cut into small discs (~5mm ×5mm). To promote the penetration of cordycepin, vacuum infiltration was performed 3 times, for 1 minute each. The *Arabidopsis* seedlings or tobacco leaf discs incubated with cordycepin were placed either at 22°C or 4°C, and harvested using a time series of 15, 30, 60, 120 and 240 minutes. For cold treatment experiments, buffer was pre-cooled at 4°C, vacuum infiltration was performed in a cold room at 4 °C. The plants without cordycepin incubation were harvested as the control, corresponding to 0 minutes of cordycepin incubation. RNA was extracted and subjected to either qRT-PCR or RNA-sequencing.

### Quantitative real-time PCR (qRT-PCR)

RNA was digested using RNase-free TURBO™ DNase (Ambion). First strand cDNA was synthesized using reverse transcriptase Superscript III (Invitrogen) and oligo dT primer with home-made RT buffer (20 mM Tris (pH 8.3), 100 mM LiCl, 3 mM MgCl_2_, 1 mM DTT). Quantitative qRT-PCR was performed with LightCycler® 480 SYBR Green I Master (Roche) using CFX96 Touch Real-Time PCR Detection System (BIORAD) according to the manufacturer’s protocol. *PP2A* (*AT1G13320*) was used as the internal control. All primers are listed in Table S7.

### Phenotype assessment

To measure primary root length, images of 1-week seedlings grown at 22°C under short-day (SD, 8/16 h, light/dark) were captured using a digital camera. The root length of more than 20 seedlings was measured using ImageJ software (NIH). For cold treatment at 4°C, plates were placed at 22°C for 2 days for germination and transferred to 4°C for further growth under the same light condition. Images were taken after 4 weeks’ growth at 4°C.

### RNA G-quadruplex structure (RG4) prediction

Plant transcriptomes were obtained from the resource data of the one thousand plant transcriptomes project (*13*). RNA G-quadruplex features were extracted according to our previous study (*8*), and further assigned to different structural subclasses of G2-RG4 (RG4 with two layers of G-quartet) or G3-RG4 (RG4 with three layers of G-quartet). The RG4 density was calculated by the count of predicted RG4s, normalized to the total amount of all four bases in the transcriptome for individual species.

### Bioclimatic variables for plants

The latitude and longitude of plant geographic distribution were obtained from observations collected by the Global Biodiversity Information Facility (https://www.gbif.org). The 433 species with over 100 observations were included in further analysis. The nineteen bioclimatic variables at representative locations were extracted from the WorldClim database (*14*). BIO1:annual mean temperature, BIO2: mean diurnal range (mean of monthly (max temp - min temp)), BIO3: isothermality (BIO2/BIO7) (×100), BIO4: temperature seasonality (standard deviation ×100), BIO5: max temperature of warmest month, BIO6: min temperature of coldest month, BIO7: temperature annual range (BIO5-BIO6), BIO8: mean temperature of wettest quarter, BIO9: mean temperature of driest quarter, BIO10: mean temperature of warmest quarter, BIO11: mean temperature of coldest quarter, BIO12: annual precipitation, BIO13: precipitation of wettest month, BIO14: precipitation of driest month, BIO15: precipitation seasonality (coefficient of variation), BIO16: precipitation of wettest quarter, BIO17: precipitation of driest quarter, BIO18: precipitation of warmest quarter, BIO19: precipitation of coldest quarter. We obtained the corresponding climate features from the latitude and longitude of each plant observation and calculated the 10, 25, 50, 75 and 90 quartiles. The Pearson correlation analysis was performed by R scripts, and the corresponding *P*-values were adjusted by Benjamini-Hochberg method (FDR). The lowest FDR among the different quartiles of climate features was used to represent the corresponding features (*15*).

### Alignment of the sequencing reads

For RNA-seq and polysome-seq libraries, reads were directly used for alignment. For SHALiPE-seq libraries, the first 3 bases at the 5’ end (random nucleotides on adaptor for cDNA ligation) of the raw reads were cropped. The alignment was carried out against *Arabidopsis* transcriptome TAIR10 release using bowtie version 1.0.1 with iterative mapping procedure (*16*). The minimum read length allowed to map was fixed to 21 bases long (*17*). The resulting mapped sam files were converted to bam files and indexed using samtools-1.4.1(*18*). The stop counts were extracted using HTseq v0.7.2 and the code was written in Python v2.7.15. Reads counts of replicates were merged after observing high correlation amongst these.

### Gini index and folding score calculation

Gin index was calculated from the SHALiPE-seq libraries with reads number of G residues in G-tract as described (*8, 19*).

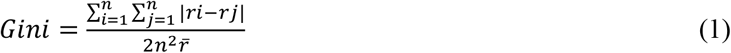

n denotes the number of G residues in the G-tracts, and ri denotes the reads number in SHALiPE profiling at position i.

To calculate the folding score, the regions with Gini (*in vitro* K^+^)/ Gini (*in vitro* Li^+^) >=1.1 and G residues with average reads count >=50 were included.

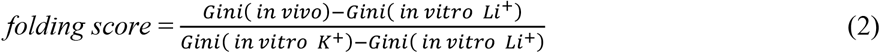

### Gene Ontology Analysis

Enrichment analysis of GO categories was performed online by AgriGo (*20*).

---

**Fig. S1.**
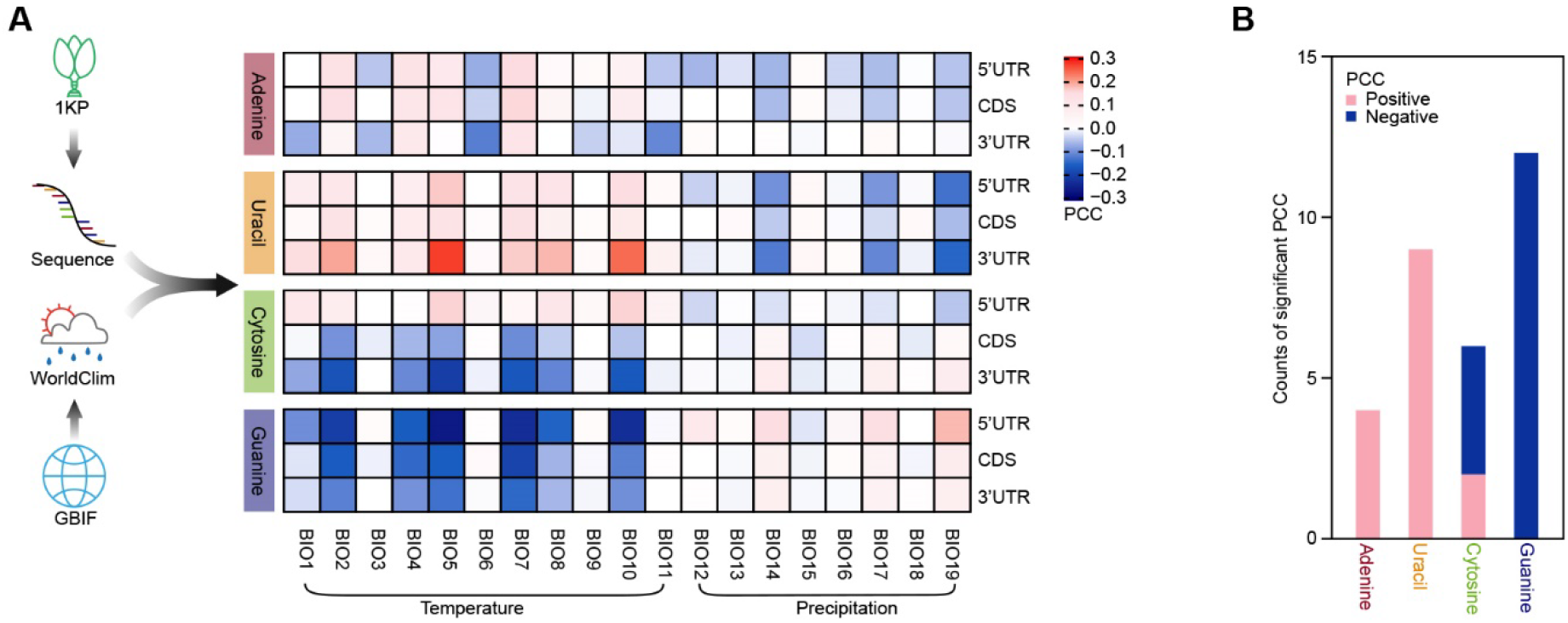
Transcriptomic nucleotide composition exhibits climatic signatures of plant adaptation. **(A)** Heat plot showing the Pearson Correlation Coefficient (PCC) between transcriptomic nucleotide frequency and bioclimatic variables of plant habitats. The frequency of the four nucleotides were calculated in 906 land plants derived from the transcriptomes of the 1000 Plants (1KP). The corresponding plant habitats were sourced from the Global Biodiversity Information Facility (GBIF). The bioclimatic variables of these plant habitats were derived from the WorldClim database (full details see Materials and Methods). BIO1: annual mean temperature; BIO2: mean diurnal range (mean of monthly (max temp - min temp)); BIO3: isothermality (BIO2/BIO7) (×100); BIO4: temperature seasonality (standard deviation ×100); BIO5: max temperature of warmest month; BIO6: min temperature of coldest month; BIO7: temperature annual range (BIO5-BIO6); BIO8: mean temperature of wettest quarter; BIO9: mean temperature of driest quarter; BIO10: mean temperature of warmest quarter; BIO11: mean temperature of coldest quarter; BIO12: annual precipitation; BIO13: precipitation of wettest month; BIO14: precipitation of driest month; BIO15: precipitation seasonality (coefficient of variation); BIO16: precipitation of wettest quarter; BIO17: precipitation of driest quarter; BIO18: precipitation of warmest quarter; BIO19: precipitation of coldest quarter, as explained by WorldClim (*https://www.worldclim.org/).* **(B)** Bar plot showing the counts of significant PCCs between transcriptomic nucleotide frequency and temperature bioclimatic variables. PCCs with a *P*-value less than a threshold of 0.01 were retained.

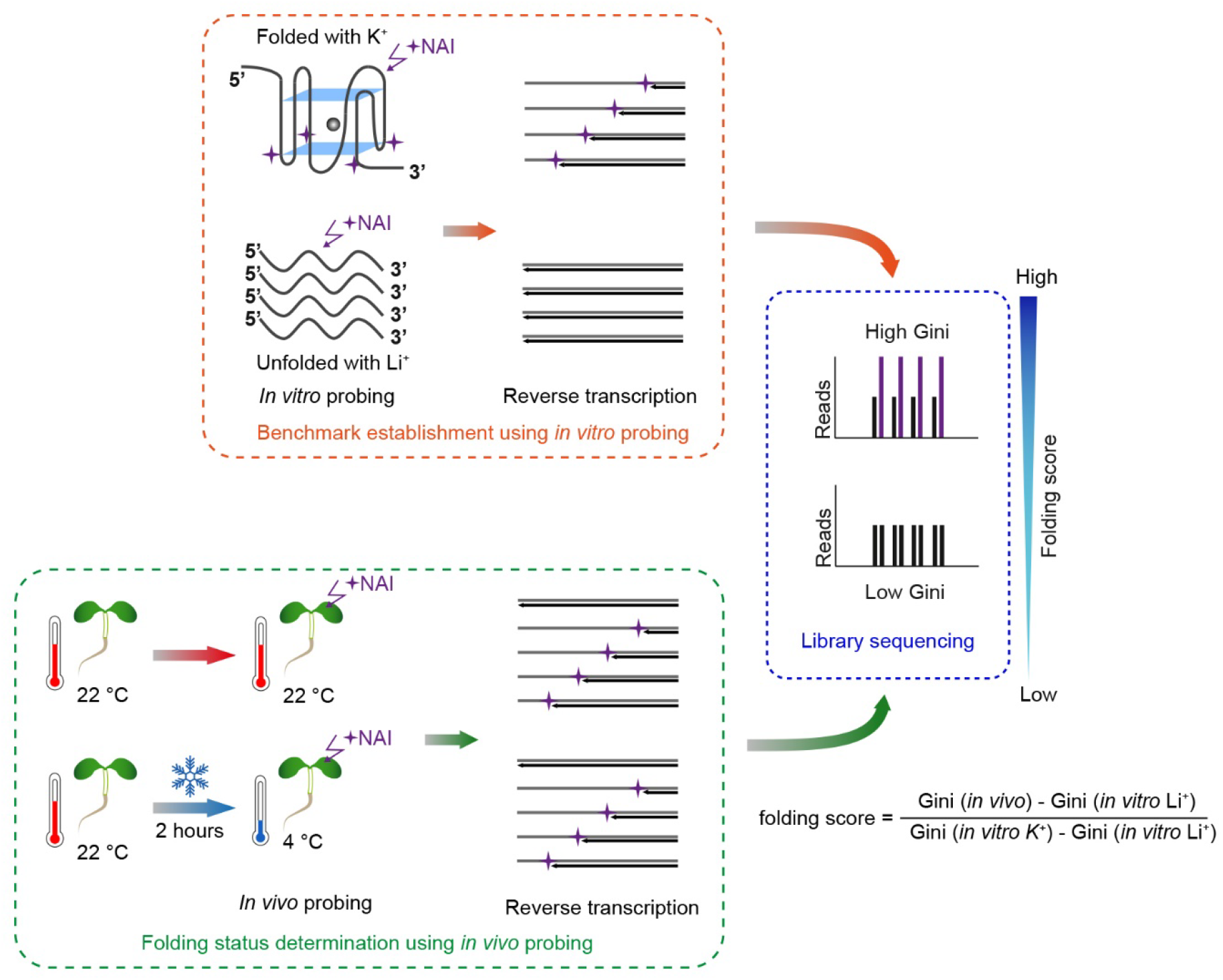
Benchmarks of folded and unfolded RNA G-quadruplex (RG4) were initially established *in vitro* (top panel) in the presence of K^+^, which stabilizes RG4 folding, or in the presence of Li^+^, which destabilizes RG4 folding. Using the SHALiPE *in vivo* probing method (lower panel), the NAI chemical (indicated by purple star) specially modifies the last G of the G-tract of folded RG4s, therefore causing high counts of reverse transcription (RT) stops with an uneven distribution, resulting in a high Gini index (middle panel on the right). While for unfolded RG4s, RT stops show an even distribution and low Gini index. For RG4 detection at 22°C and 4°C *in vivo*, NAI modification was carried out at the corresponding temperature, followed by reverse transcription, library generation and sequencing. The reads distribution and Gini index at 22°C and 4°C *in vivo* were compared to that of K^+^ *in vitro*, or Li^+^ *in vitro*, respectively.

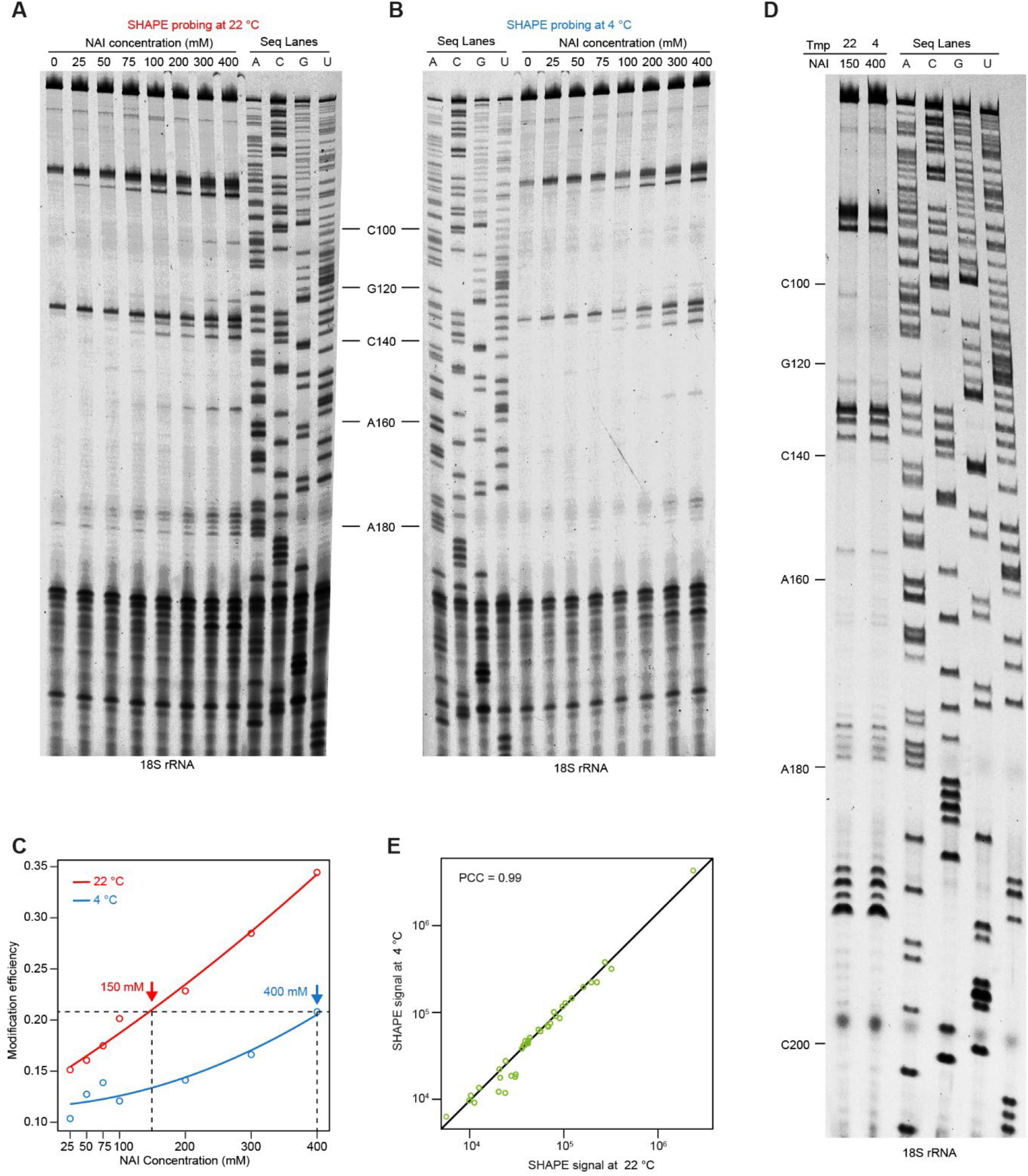
Comparable NAI modification in *Arabidopsis* at 22°C and 4°C. **(A** and **B)** Gel based analysis of 18S rRNA for chemical probing of *Arabidopsis* at 22°C and 4°C, respectively. **(C)** Modification curves of NAI on 18S rRNA at 22°C and 4°C. Modification efficiency was calculated by normalizing the signal of modified nucleotides to that of the whole lane in Fig. S3A and Fig. S3B. **(D** and **E)** Comparison of chemical probing *in vivo* at 22°C and 4°C. Gel based analysis shows comparable NAI probing profile of 18S rRNA (**D**) and high Pearson Correlation Coefficient (**E**, PCC = 0.99), with concentrations of 150mM NAI at 22°C and 400 mM NAI at 4°C, as determined in Fig. S3C.

**Fig. S4.**
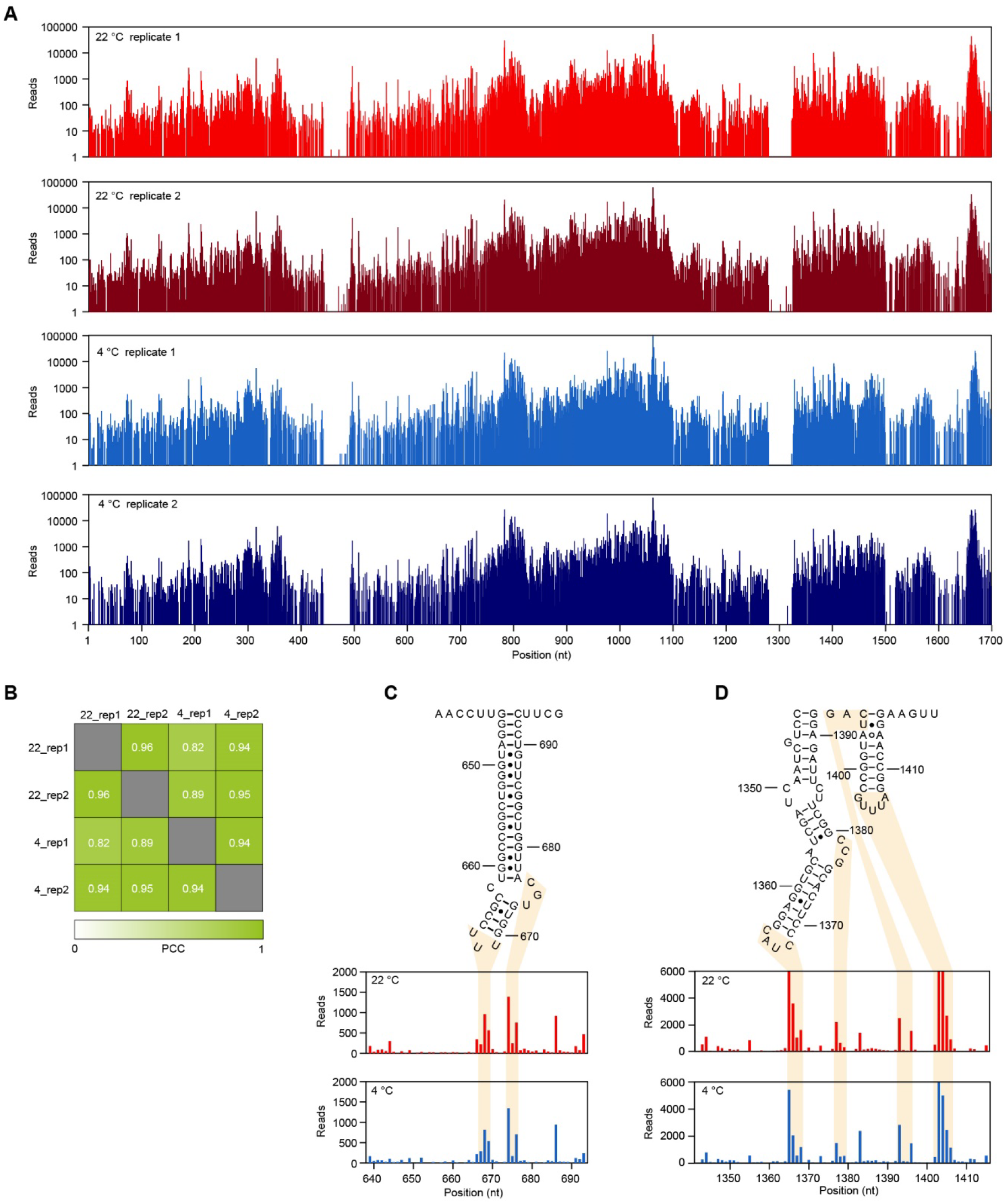
Robust and accurate determination of RNA structure using SHALiPE-seq at 22°C and 4°C. **(A)** The RT-stops of SHALiPE-seq on each nucleotide of 18S rRNA at 22°C and 4°C. **(B)** High correlations of RT-stops between independent biological replicates and different temperatures. **(C** and **D)** High agreement between SHALiPE-seq profile and phylogenetic structure on 18S rRNA. The unpaired regions in the phylogenetic structure showing high RT-stops at 22°C and 4°C are highlighted by light orange.

**Fig. S5.**
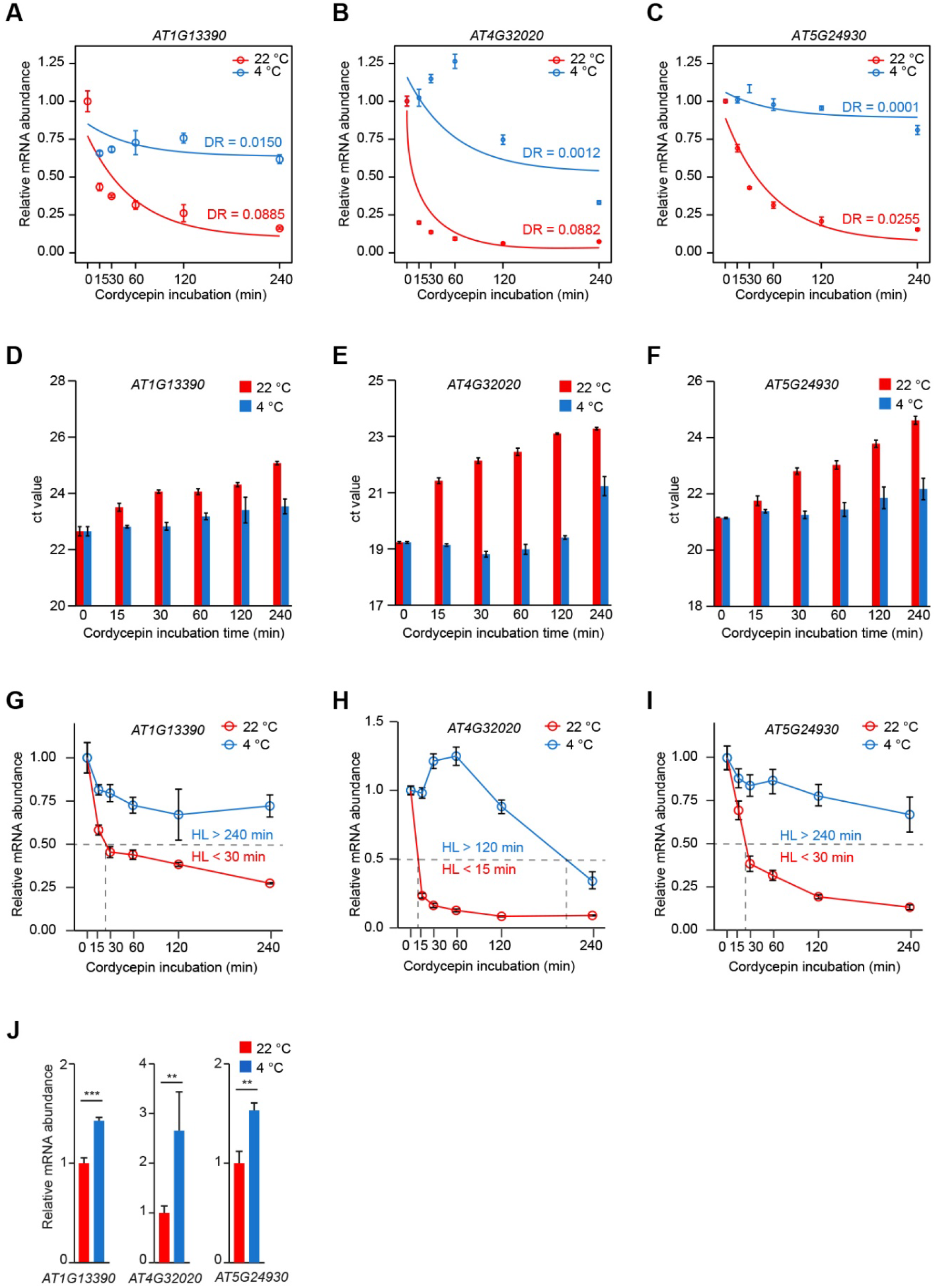
RG4 in 3’UTR represses mRNA decay and increases mRNA abundance in cold. **(A-C)** Line plot showing RNA abundance profiles with transcriptional arrest at 22°C or 4°C for selected transcripts *AT1G13390*, *AT4G32020* and *AT5G24930* with cold-responsive RG4 in 3’UTRs. For these transcripts, the decay rate (DR) was significantly smaller at 4°C compared to that at 22°C, indicating slower mRNA decay in the cold. Data were extracted from RNA-seq experiments, error bar indicates SE of 3 biological replicates. **(D-F)** Validation of RNA decay using qRT-PCR for selected transcripts at 22°C and 4°C. For *AT1G13390* **(D)**, *AT4G32020* **(E)** and *AT5G24930* **(F)**, ct values strongly increased according to transcriptional arrest duration at 22°C, suggesting a significant decrease of mRNA abundance with high decay rates at 22°C. The ct values at 4°C increased very slowly, suggesting high stability of mRNA abundance with slow decay at 4°C. **(G-I)** Line plot showing RNA abundance profiles of selected transcripts with transcriptional arrest at 22°C or 4°C. Relative mRNA abundance of *AT1G13390* **(G)**, *AT4G32020* **(H)** and *AT5G24930* **(I)** was normalized to the RNA abundance before transcriptional arrest, using the data shown in Fig. S5D-S5F. *AT1G13320 (PP2A)* was used as an internal control. HF: half-life, error bar indicates SE of 3 biological replicates. **(J)** Comparison of mRNA abundance at 22°C and 4°C for selected transcripts with cold-responsive RG4 in 3’UTR. RNA abundance was normalized to internal control of *AT1G13320*, error bars indicate SE of 3 biological replicates.

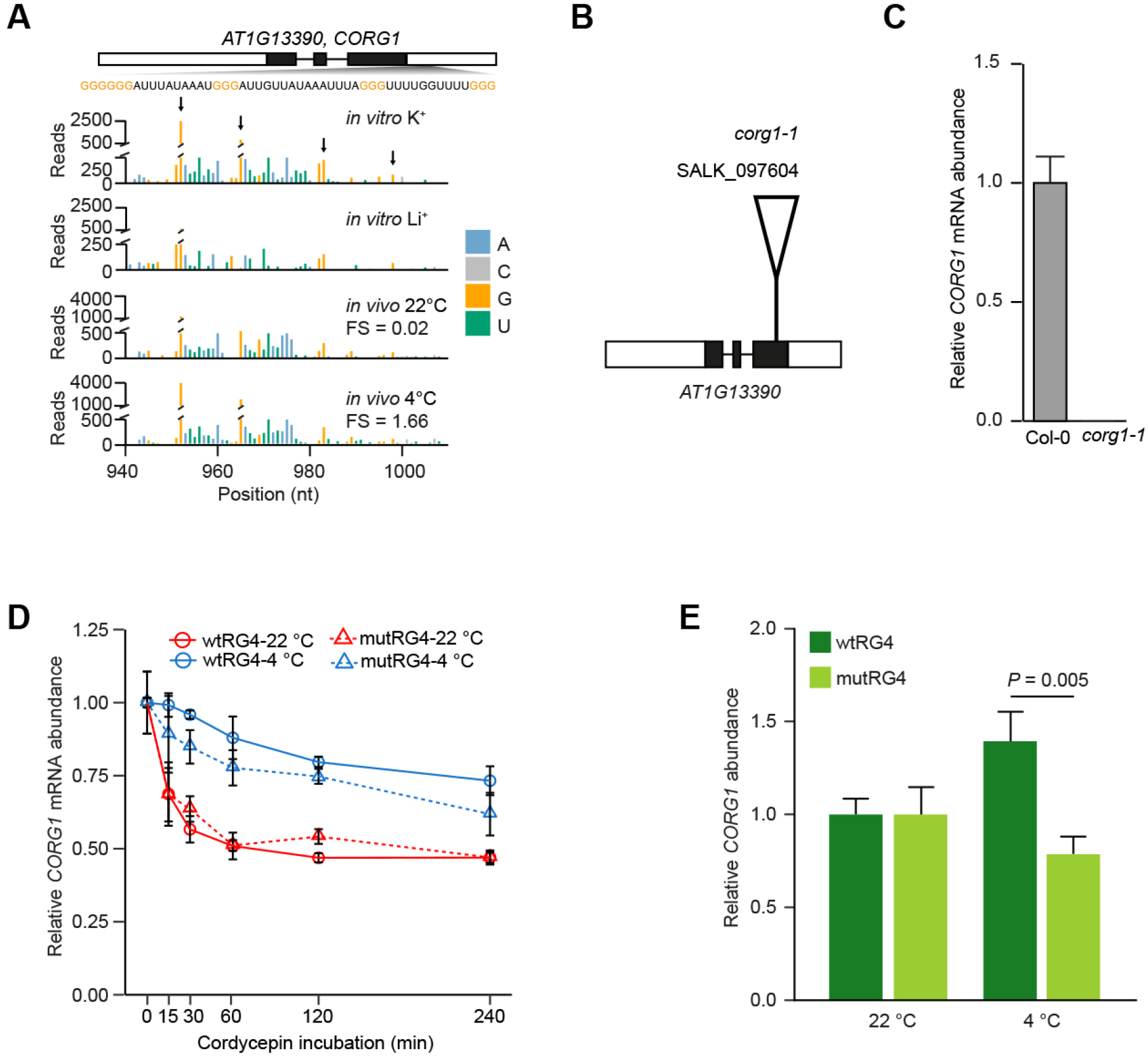
**(A)** SHALiPE-seq profiles of the RG4s in 3’UTR of *AT1G13390* (*CORG1*). High reads on last Gs (dark arrow) are shown *in vitro* with K^+^, compared to that *in vitro* with Li^+^. *In vivo* profiles show unfolded state at 22°C resembles that with Li^+^, but folded state at 4°C resembles that with K^+^. FS: folding score. **(B)** Schematic diagram of *CORG1* showing the T-DNA insertion site of *corg1-1* (SALK_097604). **(C)** Relative mRNA abundance of *CORG1* in Col-0 and *corg1-1* plants indicating *corg1-1* is a null mutant, error bar indicates SE of 3 biological replicates. **(D)** RNA decay curves at 22°C and 4°C for endogenous *CORG1* transcripts with 3’UTR carrying wtRG4 or mutRG4. Relative mRNA abundance was normalized to the RNA abundance before transcriptional arrest with cordycepin incubation, according to internal control of *PP2A*. Error bar indicates SE of 4 biological replicates. **(E)** Relative mRNA abundance of *CORG1* at 22°C and 4°C in complemented *corg1-1* mutant using genomic DNA sequence of *CORG1* carrying wtRG4 or mutRG4 in 3’UTR, respectively. Error bar indicates SE of 4 biological replicates.

